# IgGM2: An All-Atom Foundation Model for Adaptive Immune Receptor Design

**DOI:** 10.64898/2026.07.09.737510

**Authors:** Jian Ma, Fandi Wu, Lin Yao, Jing Gao, Rubo Wang, Qifeng Li, Nianzu Yang, Songlin Jiang, Dawei Huang, Xiaoyong Pan, Yiheng Zhu, Tingjun Hou, Jianhua Yao, Junchi Yan

## Abstract

Accurate immune receptor design requires modeling the coupled variation of aminoacid sequence, full-atom conformation, and target-binding geometry across antibodies, nanobodies, and T-cell receptors (TCRs). Existing methods often address only part of this problem, either by separating structure generation from sequence design, relying on fixed-backbone inverse folding, or focusing on a single receptor class. We introduce **IgGM2**, a unified all-atom generative framework for immune receptor structure prediction and CDR sequence–structure co-design. IgGM2 follows a structure-to-design strategy: it first learns how immune receptors are positioned around fixed target structures, and then transfers this target-conditioned structural prior to CDR design. Unlike modular design pipelines, IgGM2 jointly generates CDR residue identities and full-atom receptor structures, allowing framework geometry to adapt to designed CDRs without separate inverse folding or external sidechain packing. Unlike continuous residue encodings based on virtual-atom geometry, IgGM2 keeps sequence prediction explicit while using atom14 placeholders only for full-atom representation. On structure prediction benchmarks, IgGM2 better captures receptor–target spatial relationships than AlphaFold3 on FoldBench and achieves strong performance on TCR–pMHC modeling. On sequence design benchmarks, IgGM2 achieves competitive amino-acid recovery and improves Rosetta-based interface preference metrics, suggesting more favorable generated binding interfaces. These results support IgGM2 as a unified all-atom framework for adaptive immune receptor structure prediction and design.

## 1 Introduction

Antibodies and T-cell receptors (TCRs) are key molecular mediators of adaptive immunity. Both recognize cognate targets through hypervariable complementarity-determining regions (CDRs), whose sequences and conformations define binding specificity and shape the paratope–epitope interface. Accurately modeling and engineering these immune receptors is therefore central to modern therapeutic discovery, from antibody development to TCR-based cell therapy. This naturally leads to two coupled computational settings: full-atom **structure prediction** of antibody/TCR– target complexes, and target-conditioned **sequence design** of CDRs together with their binding conformations. Despite these shared molecular principles, existing methods typically study antibodies and TCRs separately, and likewise treat prediction and design as largely independent problems.

Recent advances in protein binder design have enabled target-conditioned generation of novel binding proteins [Watson et al., 2023, Bennett et al., 2026, Zhang et al., 2025]. However, many methods still follow a modular paradigm, where a binder structure is first generated and its sequence is then recovered by inverse folding [Watson et al., 2023, Dauparas et al., 2022]. Although practical, this strategy treats structure generation and sequence design as separate problems, rather than modeling their coupling directly. Recent all-atom design models attempt to unify sequence and structure generation through fixed-size atom slots and virtual atoms [Butcher et al., 2025, Qu et al., 2024, 2025], but residue identity is encoded through artificial geometric patterns. Since these virtual atoms do not correspond to real physical atoms, this representation is computationally convenient but less natural from a biochemical perspective. Antibody-specific methods introduce further assumptions: some require resolved or pre-docked antibody–antigen complexes, while others design CDRs on fixed frameworks or rely on templates, which can constrain generation in practical discovery settings [Luo et al., 2022, Kong et al., 2023, Wu et al., 2025c]. These limitations motivate a framework that can explicitly model amino acid sequences and full-atom structures together, without relying on separate inverse folding or virtual-atom residue encodings.

We introduce **IgGM2**, a unified all-atom framework for immune receptor structure prediction and target-conditioned CDR sequence–structure co-design. Our key idea is to use **structure prediction as the geometric foundation for design**. As illustrated in Figure 1a, the structure prediction model learns how antibody, nanobody, and TCR binders are positioned around fixed targets such as antigens or pMHCs. IgGM2 transfers this target-conditioned structural prior to CDR design, where residue identities and full-atom receptor conformations are generated jointly. IgGM2 differs from existing design pipelines in two main ways. First, it couples CDR sequence generation with full-atom structural denoising, **rather than generating a backbone followed by inverse folding**. Second, it uses atom14 placeholders only to maintain a fixed-size atom representation during sequence design, rather than as artificial geometric codes for residue identity. Directly predicting such virtual atoms would introduce nonphysical targets, since these atoms do not correspond to real chemical entities.

**Figure 1:**
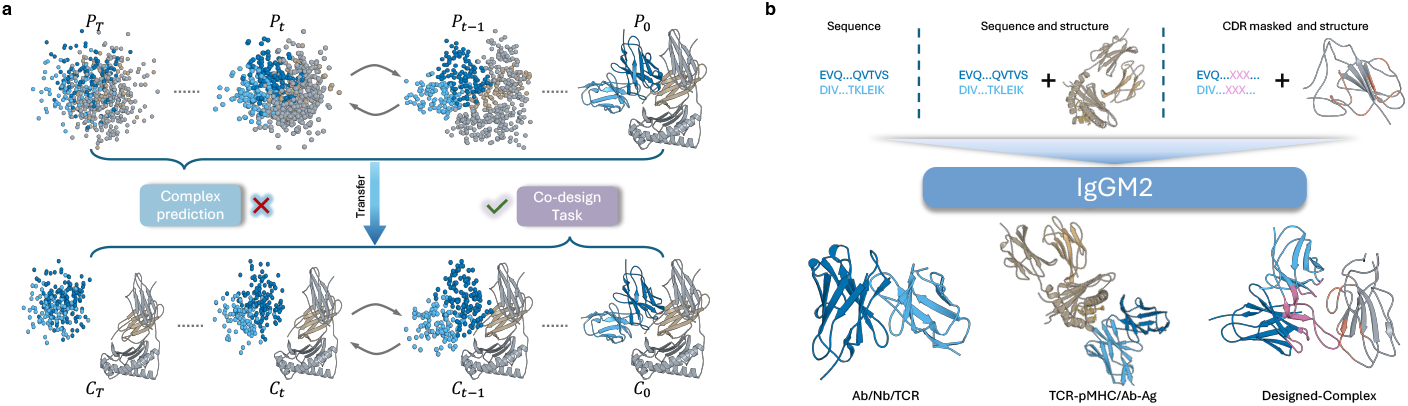
Overview of IgGM2. (a) IgGM2 performs receptor-atom diffusion conditioned on fixed structural context for both immune-complex prediction and CDR co-design. (b) It supports antibody/nanobody–antigen and TCR–pMHC complex prediction, antibody and TCR monomer prediction, and framework-conditioned CDR co-design.

As shown in Figure 1b, IgGM2 covers antibody, nanobody, TCR, and immune-complex modeling, and is instantiated as two task-specific variants: **IgGM2-P** for structure **P**rediction and **IgGM2-D** for sequence–structure co-**D**esign. Together, IgGM2 provides a structure-to-design route for immune receptor modeling, enabling full-atom CDR generation without separate inverse folding or external sidechain packing. Our main contributions are as follows:

- We develop **IgGM2**, a unified framework for antibody, nanobody, and TCR monomer prediction, immune-complex prediction, and target-conditioned receptor design.
- We propose mixed training and staged inference for CDR co-design, jointly generating CDR sequences and refining full-atom structures without external sidechain packing.
- IgGM2 achieves strong performance on structure prediction and CDR co-design benchmarks, demonstrating its effectiveness across antibody and TCR modeling tasks.

## 2 Related work

### Antibody modeling and design

Deep learning has rapidly advanced antibody structure prediction and generative design. Structure predictors such as IgFold and ABodyBuilder2 provide accurate variable-domain modeling [Ruffolo et al., 2023, Abanades et al., 2023], while generative models such as DiffAb and dyMEAN introduced antigen-conditioned CDR design [Luo et al., 2022, Kong et al., 2023, Yang et al., 2025]. Later frameworks such as FlowDesign and AbFlow improved sequence-structure co-design [Wu et al., 2025c, Wang et al., 2026]. More recent systems, such as Latent-X2 and Origin-1, have moved toward full-atom, developability-aware antibody design through increasingly integrated design pipelines [Latent Labs Team et al., 2025, Levine et al., 2026]. Despite this progress, most existing methods remain focused on antibodies alone rather than broader immune receptor modeling.

### TCR modeling and design

TCR modeling has also progressed, though it remains less developed than antibody design. Models such as TCRModel2 and TCRBuilder2 improve structural prediction [Yin et al., 2023, Abanades et al., 2023], while TCR-VALID, TCR-TRANSLATE, TCRdesign, and TCR-epiDiff explore sequence generation and epitope-conditioned design [Leary et al., 2024, Karthikeyan et al., 2025, Li et al., 2025, Seo and Rhee, 2025]. However, most current TCR approaches still focus primarily on sequence-level generation or partial structural design, with relatively few unified full-atom frameworks.

### Modular and general binder design

General binder design methods commonly rely on either modular or integrated pipelines. Two-stage approaches such as RFdiffusion combined with ProteinMPNN first generate structures and then recover sequences [Watson et al., 2023, Dauparas et al., 2022], a strategy also used in RFantibody [Bennett et al., 2026]. More recent frameworks, including ODesign and PPIFlow, move toward more integrated all-atom generation while still often incorporating inverse folding, refinement, or affinity maturation stages [Zhang et al., 2025, Yu et al., 2026]. These methods have substantially expanded binder design capabilities, but sequence and structure generation are often still partially separated.

## 3 Method

### 3.1 Overview

IgGM2 follows a structure-to-design strategy. For immune receptor modeling, the model must first learn how antibodies, nanobodies, and TCRs are positioned relative to molecular targets. We therefore train **IgGM2-P** as a context-conditioned structure prediction model. For monomer prediction, IgGM2-P generates the full immune receptor structure from sequence. For complex prediction, the antigen or pMHC is fixed as the target context, and the immune receptor structure is generated around it. **IgGM2-D** transfers this structural prior to target-conditioned CDR co-design. In this setting, the target structure is fixed, and non-designed receptor residues keep their original amino acid identities, while their coordinates are still generated together with the designed CDRs. This allows the framework geometry to adapt to CDR sequence and conformational changes, rather than being treated as a rigid scaffold. IgGM2-D couples sequence and structure generation by aligning discrete sequence masking with continuous EDM coordinate noise, and uses staged inference to first resolve CDR sequence and backbone geometry before refining the full-atom structure.

We organize the method into four parts: EDM coordinate diffusion, noise-aligned sequence masking, IgGM2-P for context-conditioned structure prediction, and IgGM2-D for full-atom CDR sequence– structure co-design with mixed training and staged sampling.

### 3.2 Preliminary: EDM Preconditioning

For coordinate diffusion, we adopt the EDM parameterization [Karras et al., 2022]. Given clean atom coordinates ***x***_0_ ∈ ℝ ^*N ×*3^, noisy coordinates at noise level σ are defined as

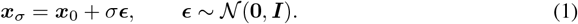

The denoiser is parameterized as

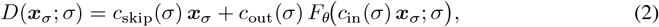

where c_in_(σ), c_out_(σ), and c_skip_(σ) follow the standard EDM formulation. In particular,

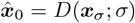

denotes the denoised estimate of the clean coordinates used for coordinate supervision and sampling updates.

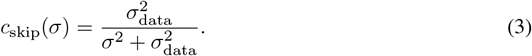

We regard c_skip_(σ) as a coordinate-side information level, which provides a continuous schedule for matching sequence visibility to structure denoising.

### 3.3 Context-Conditioned Structure Prediction

We build IgGM2-P on an AF3-like diffusion architecture for immune receptor structure prediction [Abramson et al., 2024]. Unlike AlphaFold-style models that rely on MSA and template inputs, we remove both modules and use a single-sequence structure model (as shown in Figure 2a), following recent observations that MSA provides limited additional benefit for antibody and TCR modeling. Since our goal is to transfer a strong structure prediction model toward immune receptor co-design, we initialize the shared AF3-like modules from pretrained Protenix parameters [ByteDance Research, 2024] and fine-tune them on immune receptor data. To support target-conditioned prediction, we further incorporate a structural constraint embedding following ODesign [Zhang et al., 2025].

**Figure 2:**
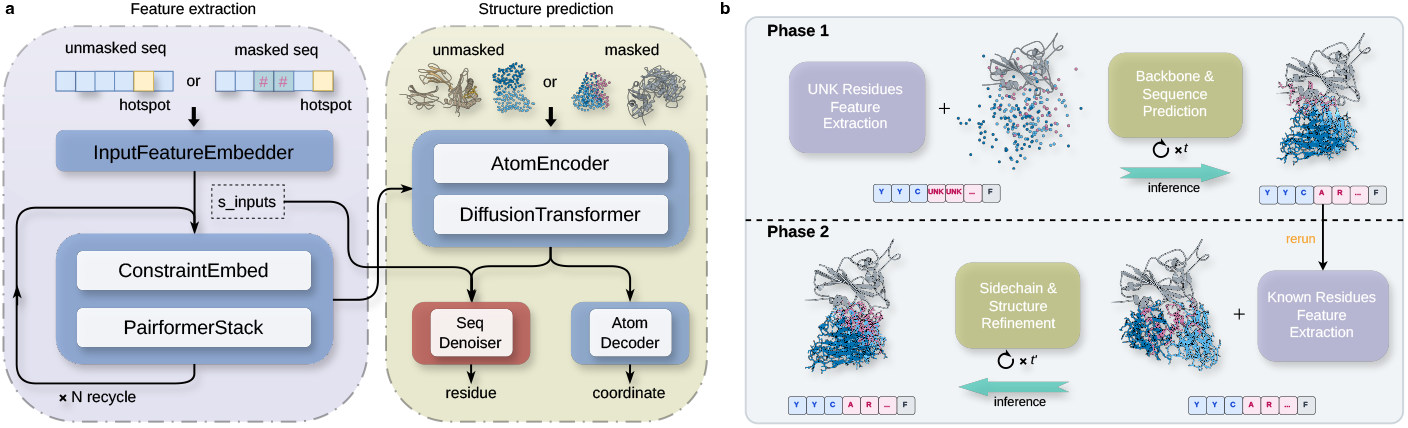
IgGM2 architecture. **(a)** The structure model injects fixed-context information into the Pairformer trunk and denoises only generated atoms. IgGM2-P uses this architecture without the sequence denoiser, while IgGM2-D adds the orange sequence-denoiser module for CDR sequence– structure co-design. **(b)** For CDR co-design, IgGM2-D uses a two-phase sampler: it first predicts CDR sequence and backbone geometry with masked CDR identities, then fixes the predicted sequence and refines the full-atom structure.

IgGM2-P supports monomer and complex structure prediction. For monomer prediction, including antibody, nanobody, and TCR modeling, no structural context is provided, and all receptor atoms are generated by diffusion. For complex prediction, such as antibody–antigen or TCR–pMHC modeling, the target structure is kept fixed, and only the receptor atoms are diffused and denoised. In target-conditioned settings, hotspot residues can also be provided to guide interface modeling.

The structural constraint embedding provides the network with information about fixed structural contexts. Fixed atoms are marked by binary masks, and their geometric features are projected into the pair representation. Hotspot residues are injected through the same conditioning pathway as token-level features. This allows the Pairformer trunk to use the fixed-context information, while the diffusion module updates only the generated atoms.

During training, coordinate supervision is applied only to generated atoms. Let m_gen_ denote the atom-level mask of generated regions. The coordinate loss is

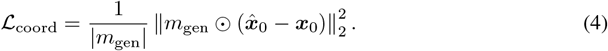

For monomer prediction, m_gen_ covers all atoms. For target-conditioned complex prediction, atoms in the fixed context are excluded from the coordinate loss. The full structure prediction objective is

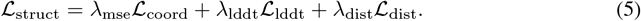

#### Algorithm 1

IgGM2-D Phase-Split Sampling

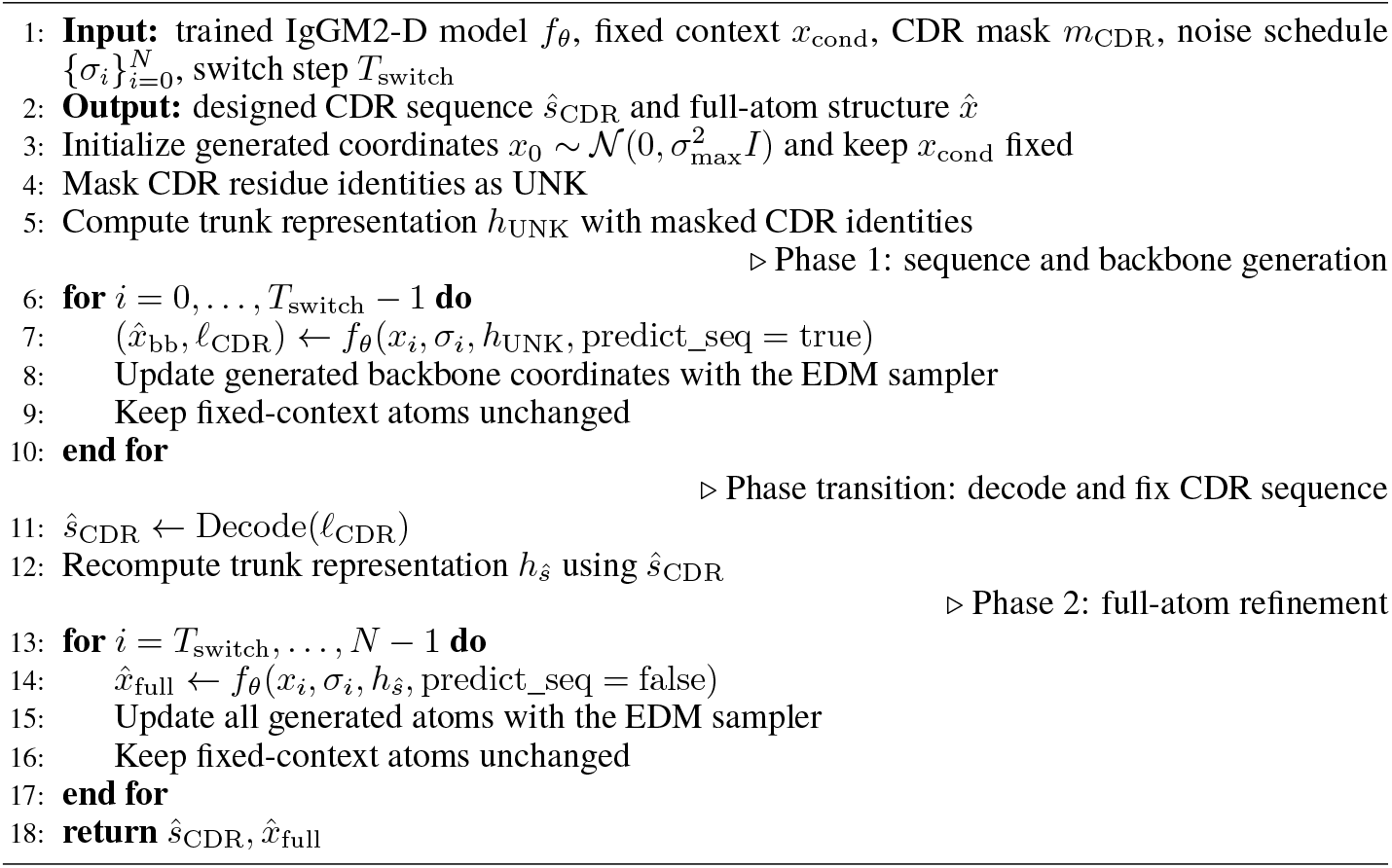

### 3.4 Sequence Mask Rate Alignment

The EDM preconditioning in Section 3.2 provides a continuous notion of coordinate reliability through c_skip_(σ). We use this quantity to define a matched corruption schedule for discrete sequence tokens. We couple discrete sequence corruption to the same diffusion noise level σ. Let p(σ) denote the masking probability for sequence tokens, such that a fraction 1 − p(σ) remains visible and the remaining tokens are masked for conditional prediction.

To maintain consistent information exposure across sequence and structure, we align the visible sequence fraction with this coordinate reliability:

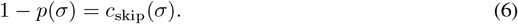

This yields

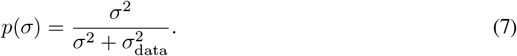

Under this schedule, low structural noise corresponds to minimal sequence masking, while high structural noise progressively increases masking toward fully de novo sequence generation. This defines a shared information-exposure schedule for sequence and structure, rather than assuming that sequence masking follows the same physical corruption process as coordinate diffusion. A more detailed signal-to-noise interpretation of this alignment is provided in Appendix A.2.

### 3.5 CDR Sequence–Structure Co-Design

We extend the structure prediction model into **IgGM2-D**, a target-conditioned CDR sequence– structure co-design model. IgGM2-D is warm-started from the trained IgGM2-P model, allowing it to inherit a strong structural prior before learning the sequence design objective. As shown in Figure 2a, we add a sequence denoiser on top of the shared trunk to predict the residue identities of designed CDR positions.

#### Mixed training

To enable sequence design while preserving full-atom structural modeling, we train IgGM2-D with a mixed strategy. For each training sample, we select one of two modes according to a mixing rate ρ. In sequence-design mode, CDR residue identities are masked using the noise-aligned schedule in Eq. 7, and the model predicts both CDR sequences and backbone structures.

In structure-prediction mode, the ground-truth CDR sequence is provided, and the model predicts full-atom structures, including sidechains, without applying a sequence loss. The training objective is

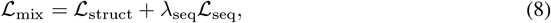

where *L*_seq_ is applied only in sequence-design mode, and *L*_struct_ follows the structure loss defined in Section 3.3. In sequence-design mode, the CDR residue identities are unknown, so structural supervision mainly focuses on CDR backbone atoms together with sequence recovery. In structure-prediction mode, the ground-truth CDR sequence is given, allowing full-atom supervision over both backbone and sidechain coordinates. This mixed training exposes IgGM2-D to both design and reconstruction signals, enabling it to learn CDR sequence generation while preserving accurate full-atom structural modeling.

#### Staged inference

At inference time, IgGM2-D uses a staged sampling procedure controlled by a noise threshold σ_switch_, as shown in Figure 2b. In the high-noise stage, CDR identities are treated as unknown, and the model jointly predicts CDR residue types and backbone geometry. When the noise level reaches σ_switch_, the predicted CDR sequence is decoded and fixed. The model then continues denoising in the low-noise stage, using the fixed sequence to refine the full-atom structure, including CDR sidechains. This staged procedure separates sequence determination from final full-atom refinement, enabling IgGM2-D to generate CDR sequences and all-atom structures without a separate inverse-folding model or external sidechain-packing tool. Detailed training and sampling procedures are provided in Algorithms 1 and S1.

## 4 Experiment

In this section, we evaluate IgGM2 on three tasks: (i) immune-complex structure prediction for antibody–antigen and TCR–pMHC systems, (ii) CDR sequence–structure co-design for paired antibodies and nanobodies, and (iii) ablation studies on the mixed-training rate and phase-transition step used in our co-design protocol. We introduce the datasets, metrics, and baselines in Section 4.1 and use the same definitions throughout all experiments.

### 4.1 Setup

#### Data

Training data are curated from SAbDab [Dunbar et al., 2014] and STCRDab [Leem et al., 2018]. Since our study includes both structure prediction benchmarking against models such as AlphaFold-3 and Chai-1 [Abramson et al., 2024, Chai Discovery, 2024] and sequence design experiments, we construct two separate datasets for these tasks. Both datasets are first deduplicated at the sequence level and then clustered by CDRH3 using MMseqs with <monospace>min_seq_id = 0.8</monospace> and cov_mode = 1. For structure prediction, the training set includes structures released before September 30, 2021. To ensure fair evaluation, we further remove training samples with full-sequence similarity above 95% to any sample in FoldBench [Xu et al., 2025] or the other test benchmarks. For sequence design, we retain only complexes with receptor–target information, as the task is to design binders under antigen-conditioned settings. We use January 1, 2024 as the temporal cutoff, assigning earlier samples to training and later samples to validation and test sets. We also apply CDRH3-based cluster splitting to ensure that CDRH3 sequence similarity between training and test sets is below 0.8.

#### Benchmarks

We evaluate on three structure prediction benchmarks in the main text (FoldBench [Xu et al., 2025], SAbDab-22H2-AbAg [Dunbar et al., 2014], STCRDab-22-TCR_pMHC [Leem et al., 2018]) and three supplementary benchmarks reported in Appendix C (SAbDab-22H2-Ab, SAbDab-22H2-Nano, STCRDab-22-TCR). For sequence design, the full clean SAbDab test split contains 269 complexes (`indices_design_sab_test_clean.csv:` 195 paired antibodies and 74 nanobodies). The train–test cluster split controls similarity between training and test complexes, but it does not remove redundant samples within the held-out test split. Because the full test set still contains many mutually similar CDRH3 sequences, the main sequence-design tables report a test-internally deduplicated subset with pairwise CDRH3 similarity below 0.8 (203 complexes; 148 paired antibodies and 55 nanobodies), while full-set results are provided in Appendix D.

#### Metrics

Structure prediction is evaluated using DockQ, success rate (SR; DockQ ≥ 0.23) and TM-Score. For sequence design, the main tables report CDR amino-acid recovery (AAR), local CDR Cα RMSD, and Rosetta interface-energy metrics, including IMP and dG_separated_ (ref2015).

Co-generated DockQ / TM-Score / Cα RMSD / SR metrics on the full clean test set are reported in Appendix D. Backbone-only design baselines are first side-chain-packed with Rosetta PackRotamersMover and then locally relaxed before IMP is computed; full-atom outputs go directly into local FastRelax.

#### Baselines

We consider a broad range of prior methods for structure prediction (AlphaFold-3 / -Multimer, Protenix, Boltz-1, HelixFold-3, Chai-1, OpenFold-3, tFold-{Ab, Ag, TCR}, TCRModel2, IgFold, ImmuneBuilder, ESMFold, OmegaFold, EquiFold, HelixFold-Single) [Abramson et al., 2024, Evans et al., 2022, ByteDance Research, 2024, Wohlwend et al., 2024, Liu et al., 2024, Chai Discovery, 2024, OpenFold Consortium, 2024, Wu et al., 2025a,b, Yin et al., 2023, Ruffolo et al., 2023, Abanades et al., 2023, Lin et al., 2023, Wu et al., 2022, Lee et al., 2022, Fang et al., 2022] and for CDR co-design (DiffAb, dyMEAN, AbX, RFAntibody, IgGM) [Luo et al., 2022, Kong et al., 2023, Zhu et al., 2024, Bennett et al., 2026, Wang et al., 2025]. Since these models are not all evaluated on the same benchmarks, we do not enforce a unified baseline set across all experiments; instead, each benchmark is compared against the methods with publicly reported results available, following the evaluation protocol of the corresponding prior work. For CDR co-design, baselines re-trained from scratch on the training data are marked with ^*∗*^; baselines without ^*∗*^ use their released checkpoints.

### 4.2 Structure Prediction

We evaluate IgGM2-P as a binder-only structure predictor on two main immune-complex benchmarks: Fold-Bench for antibody/nanobody–antigen complexes and STCRDab-22-TCR_pMHC for TCR–pMHC complexes. IgGM2-P uses a single 1×1 sample without candidate selection across both benchmarks. Additional results on SAbDab-22H2-AbAg complex prediction and separate antibody, nanobody, and TCR monomer prediction benchmarks are provided in Appendix C.

#### Antibody/nanobody–antigen complex prediction

On FoldBench, IgGM2-P achieves a strong success rate of 73.8% using only a single 1 × 1 sample without candidate ranking, outperforming the official leaderboard baselines (Table 1). This setting is also more efficient: the compared baselines follow a 5 × 5 sampling protocol, generating multiple candidates across five seeds and selecting the best-ranked prediction, whereas IgGM2-P produces one prediction directly without post-hoc selection. These results support the effectiveness of our binder-only formulation when the antigen is available as a fixed structural context. Representative examples further show that IgGM2-P preserves antibody/nanobody–antigen interface geometry across both receptor types (Figure 3). For the 8ATH antibody–antigen complex, IgGM2-P improves DockQ from 0.286 to 0.721 and reduces global RMSD from 4.16Å to 1.50Å compared with AlphaFold-3. For the 8AOK nanobody–antigen complex, it improves DockQ from 0.507 to 0.577 and reduces global RMSD from 2.58Å to 1.87Å. These examples indicate that fixing the antigen context helps IgGM2-P accurately place and model immune receptor binders.

**Table 1:** FoldBench antibody-antigen prediction. Baselines follow the official leaderboard settings, while IgGM2-P uses a single 1 × 1 sample without candidate ranking because it has no confidence head.

| Method | SR (%) ↑ | Mean DockQ ↑ |
| --- | --- | --- |
| Chai-1 | 23.6 | 0.17 |
| HelixFold-3 | 28.4 | 0.21 |
| Boltz-1 | 33.5 | 0.22 |
| Protenix-0.5 | 34.1 | 0.23 |
| AlphaFold-3 | 47.9 | 0.36 |
| Protenix-1.0 | 48.8 | 0.36 |
| **IgGM2-P(epitope)** | **73.8** | **0.60** |

**Figure 3:**
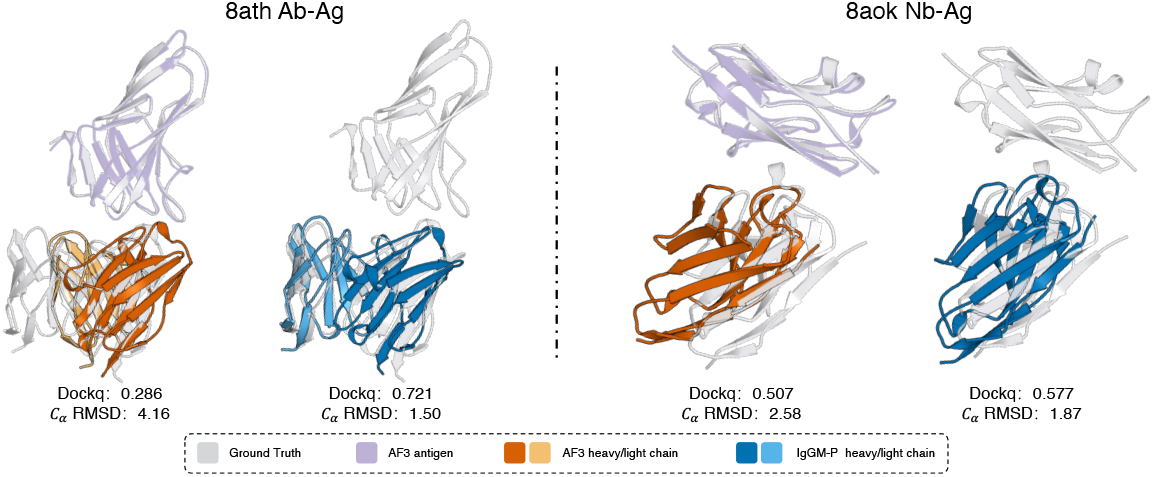
FoldBench predictions. Left: 8ATH antibody–antigen complex. Right: 8AOK nanobody– antigen complex.

Since IgGM2-P conditions on fixed target structures and can incorporate epitope-level guidance, we further evaluate it on SAbDab-22H2-AbAg against methods that use comparable contact or epitope restraints. As shown in Table S2, IgGM2-P outperforms existing epitope-conditioned baselines, demonstrating that epitope-guided structural constraints provide effective interface guidance.

#### STCRDab-22-TCR_pMHC complex prediction

TCR–pMHC prediction is particularly challenging because the complex contains multiple interacting components, including the TCR, peptide, and MHC molecules. Rather than predicting the full complex from scratch, IgGM2-P keeps the pMHC structure fixed and diffuses only the TCR binder. As shown in Table 2, this formulation achieves the best overall performance, with DockQ 0.746, full-complex GDT 0.887, and acceptable/medium/high success rates of 100.0/88.9/38.9%. Additional prediction examples are provided in Appendix F.1. These results show that binder-only diffusion is effective for TCR–pMHC modeling, where the peptide–MHC structure provides a strong conditioning context.

**Table 2:** TCR–pMHC complex prediction on STCRDab-22-TCR_pMHC (18 complexes). Baselines use official FASTA/native-PDB inputs. Acc., Med., and High SR are DockQ success rates at thresholds 0.23, 0.49, and 0.80.

| Method | DockQ ↑ | LRMS ($\text{\AA}$) ↓ | iRMS ($\text{\AA}$) ↓ | GDT ↑ | Acc. SR (%) ↑ | Med. SR (%) ↑ | High SR (%) ↑ |
| --- | --- | --- | --- | --- | --- | --- | --- |
| TCRModel2 | 0.585 | 7.749 | 2.932 | 0.795 | 88.9 | 66.7 | 16.7 |
| AlphaFold-3 | 0.632 | 5.316 | 2.017 | 0.816 | 94.4 | 77.8 | 33.3 |
| tFold-TCR | 0.523 | 5.585 | 2.242 | 0.774 | 100.0 | 55.6 | 5.6 |
| AlphaFold-Multimer | 0.537 | 6.052 | 2.293 | 0.775 | 100.0 | 44.4 | 16.7 |
| **IgGM2-P(epitope)** | **0.746** | **2.966** | **1.262** | **0.887** | **100.0** | **88.9** | **38.9** |

### 4.3 CDR Sequence–Structure Co-Design

Following the evaluation setup in Section 4.1, we report the main sequence-design results on the CDRH3-deduplicated SAbDab test subset. This subset includes paired antibodies and nanobodies, with CDR recovery and local structural accuracy reported in Tables 3 and 4. We further report Rosetta-based interface metrics in Table 5. Full-set results and additional complex-quality metrics are provided in Appendices D.2 and D. TCR CDR design is not included in the main evaluation; we provide an additional all-six-CDR TCR design study in Appendix D.3.

**Table 3:**
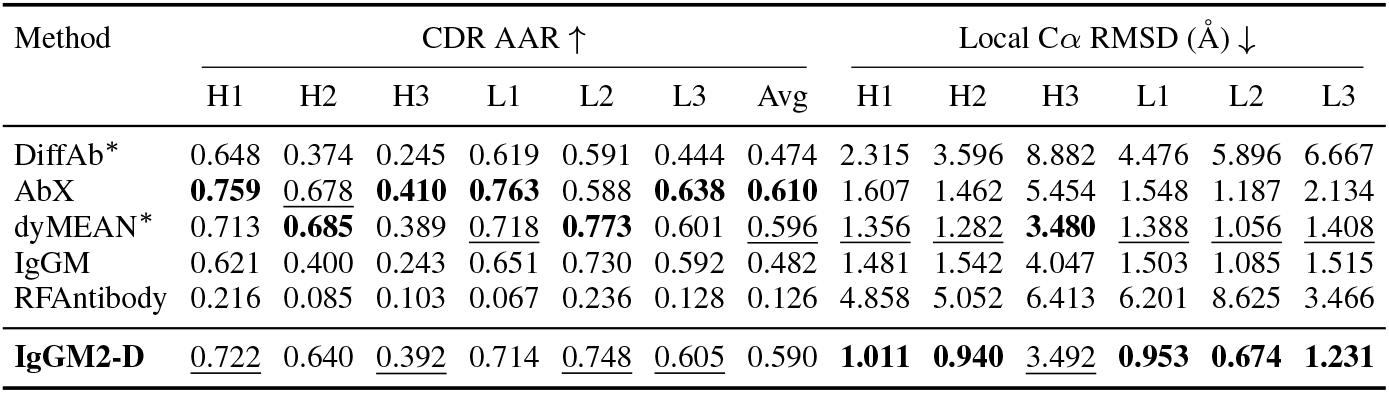
Paired-antibody CDR co-design performance on the CDRH3-deduplicated SAbDab design test subset. Higher AAR and lower framework-aligned local Cα RMSD indicate better performance. Best values are bolded and second-best values are underlined. HDOCK is used for DiffAb and AbX when external docking is required.

| Method | CDR AAR ↑ | | | | | | | Local $C\alpha$ RMSD ($\text{\AA}$) ↓ | | | | | |
| --- | --- | --- | --- | --- | --- | --- | --- | --- | --- | --- | --- | --- | --- |
|  | H1 | H2 | H3 | L1 | L2 | L3 | Avg | H1 | H2 | H3 | L1 | L2 | L3 |
| DiffAb* | 0.648 | 0.374 | 0.245 | 0.619 | 0.591 | 0.444 | 0.474 | 2.315 | 3.596 | 8.882 | 4.476 | 5.896 | 6.667 |
| AbX | **0.759** | <u>0.678</u> | **0.410** | **0.763** | 0.588 | **0.638** | **0.610** | 1.607 | 1.462 | 5.454 | 1.548 | 1.187 | 2.134 |
| dyMEAN* | 0.713 | **0.685** | 0.389 | <u>0.718</u> | **0.773** | 0.601 | <u>0.596</u> | <u>1.356</u> | <u>1.282</u> | **3.480** | <u>1.388</u> | <u>1.056</u> | <u>1.408</u> |
| IgGM | 0.621 | 0.400 | 0.243 | 0.651 | 0.730 | 0.592 | 0.482 | 1.481 | 1.542 | 4.047 | 1.503 | 1.085 | 1.515 |
| RFAntibody | 0.216 | 0.085 | 0.103 | 0.067 | 0.236 | 0.128 | 0.126 | 4.858 | 5.052 | 6.413 | 6.201 | 8.625 | 3.466 |
| **IgGM2-D** | <u>0.722</u> | 0.640 | <u>0.392</u> | 0.714 | <u>0.748</u> | <u>0.605</u> | 0.590 | **1.011** | **0.940** | <u>3.492</u> | **0.953** | **0.674** | **1.231** |

**Table 4:** Nanobody CDR co-design performance on the CDRH3-deduplicated SAbDab design test subset. AAR is higher better; framework-aligned local Cα RMSD is lower better.

| Method | CDR AAR ↑ | | | | Local $C\alpha$ RMSD ($\text{\AA}$) ↓ | | |
| --- | --- | --- | --- | --- | --- | --- | --- |
|  | H1 | H2 | H3 | Avg | H1 | H2 | H3 |
| DiffAb* | <u>0.487</u> | <u>0.335</u> | <u>0.177</u> | <u>0.287</u> | <u>2.506</u> | 1.794 | 4.992 |
| IgGM | 0.468 | 0.291 | 0.154 | 0.267 | 2.679 | 2.311 | 5.020 |
| RFAntibody | 0.181 | 0.097 | 0.115 | 0.127 | 5.092 | 4.391 | 7.445 |
| **IgGM2-D** | **0.497** | **0.499** | **0.238** | **0.372** | **2.328** | **1.597** | **4.374** |

**Table 5:** Rosetta interface-energy metrics for CDR co-design on the CDRH3-deduplicated SAbDab design test subset.

| Method | Paired antibodies |  | Nanobodies |  |
| --- | --- | --- | --- | --- |
| | IMP ↑ | Mean $dG$ ↓ | IMP ↑ | Mean $dG$ ↓ |
| DiffAb* | 0.392 | 29.86 | 0.236 | 10.52 |
| AbX | 0.318 | 2.70 | — | — |
| dyMEAN* | 0.327 | 2191.46 | — | — |
| IgGM | <u>0.541</u> | -17.83 | 0.273 | <u>-9.46</u> |
| RFAntibody | 0.466 | 118.90 | <u>0.327</u> | 1.25 |
| **IgGM2-D** | **0.615** | **-23.27** | **0.527** | **-24.84** |

#### Paired antibody design performance

We first evaluate IgGM2-D on paired antibody CDR design, considering both sequence recovery and generated structural quality (Table 3). IgGM2-D does not always achieve the highest AAR, as AbX and dyMEAN recover more residues on some CDRs. However, AAR should be interpreted together with the design setting and the generated structure quality. AbX requires a completed antibody structure and external docking in our evaluation pipeline, rather than generating the bound complex end to end. It also mainly edits CDR residues while keeping the framework conformation fixed, which can be restrictive because CDR mutations may require framework-level adjustment to support the final binding geometry. IgGM shows lower AAR in our split, partly because many test antigens contain multiple chains, whereas IgGM was originally designed for single-chain antigen inputs. Although these antigens are merged into a single chain during evaluation, this input mismatch may still affect its performance. dyMEAN can generate antibody structures, but its interface-energy results are much weaker than IgGM2-D. As shown in Table 5, IgGM2-D achieves the best IMP and a mean dG of −23.27, suggesting more favorable binding geometries. It also obtains the best local Cα RMSD on five of six paired-antibody CDRs, indicating that its sequence designs are supported by more coherent local conformations.

#### Nanobody design performance

We next evaluate IgGM2-D on nanobody CDR co-design, as shown in Table 4. Nanobody design is more challenging because nanobodies usually contain a longer HCDR3 loop, making sequence recovery and local structure generation harder. IgGM2-D achieves the best average AAR, improving from 0.287 for the strongest baseline DiffAb to 0.372. On HCDR3, IgGM2-D improves AAR from 0.154 for IgGM to 0.238, corresponding to a 54.55% relative gain. It also achieves the best local Cα RMSD on all three nanobody CDRs. Consistently, Table 5 shows that IgGM2-D improves nanobody IMP from 0.327 to 0.527 and reaches a mean dG of −24.84, suggesting more favorable generated interfaces.

### 4.4 Ablations

We ablate the two main hyperparameters in the IgGM2-D co-design protocol: the mixed-training ratio between sequence-design and structure-prediction samples, and the phase-transition step used during sampling. In the main text, we analyze these factors separately, with the full grid including the backbone-noise setting reported in Appendix E.1.

#### Phase-transition step

We first study the phase-transition step under a 30/70 mixed-training setting, where structure-prediction samples are used more frequently to stabilize full-atom generation. With this ratio fixed, we vary T_switch_ to determine when the sampler should switch from sequence-and-backbone generation to full-atom refinement. We select the transition step mainly by All AAR, which measures overall sequence recovery across paired antibody and nanobody designs. As shown in Table 6, T_switch_=75 achieves the highest All AAR of 0.530, although it does not give the best IMP.

**Table 6:** Effect of the phase-transition step with seq:struct ratio 30/70 and backbone-noise scale 0.1. Metrics are rows and transition steps are columns; the selected setting is bolded.

| Metric | 25 | 50 | **75** | 200 |
| --- | --- | --- | --- | --- |
| All AAR ↑ | 0.523 | 0.528 | **0.530** | 0.522 |
| Paired AAR ↑ | 0.579 | 0.589 | **0.587** | 0.582 |
| Nano AAR ↑ | 0.373 | 0.369 | **0.379** | 0.362 |
| HCDR3 RMSD ($\text{\AA}$) ↓ | 3.846 | 3.830 | **3.702** | 3.618 |
| IMP ↑ | 0.595 | 0.561 | **0.584** | 0.543 |

#### Mixed-training ratio

After fixing T_switch_=75, we vary the fraction of sequence-design samples from 30% to 70% during mixed training (Table 7). Increasing the sequence-design ratio does not consistently improve AAR. Instead, the 30/70 setting achieves the highest All AAR and the lowest FR-aligned HCDR3 RMSD, where structures are aligned by the framework region before measuring HCDR3 deviation. This suggests that stronger structure-prediction supervision helps preserve conformational generation ability, which is important for robust sequence–structure co-design.

**Table 7:** Effect of the mixed-training ratio with backbone-noise scale 0.1 and T_switch_=75. Metrics are rows and ratios are columns; the selected setting is bolded.

| Metric | **30/70** | 50/50 | 70/30 |
| --- | --- | --- | --- |
| All AAR ↑ | **0.530** | 0.530 | 0.515 |
| Paired AAR ↑ | **0.587** | 0.589 | 0.573 |
| Nano AAR ↑ | **0.379** | 0.374 | 0.363 |
| HCDR3 RMSD ($\text{\AA}$) ↓ | **3.702** | 3.858 | 3.721 |
| IMP ↑ | **0.584** | 0.524 | 0.677 |

## 5 Discussion

IgGM2 unifies immune receptor structure prediction and CDR sequence–structure co-design within a single all-atom diffusion framework. By transferring structural modeling to target-conditioned design, IgGM2 generates CDR sequences and all-atom structures without separate inverse folding or external sidechain packing. We discuss remaining limitations, including sequence recovery and sidechain refinement, in Appendix B. Future work will explore stronger sequence priors, improved all-atom optimization, developability-aware objectives, and experimental validation.

## Appendix

### Algorithm S1

IgGM2-D Mixed Training

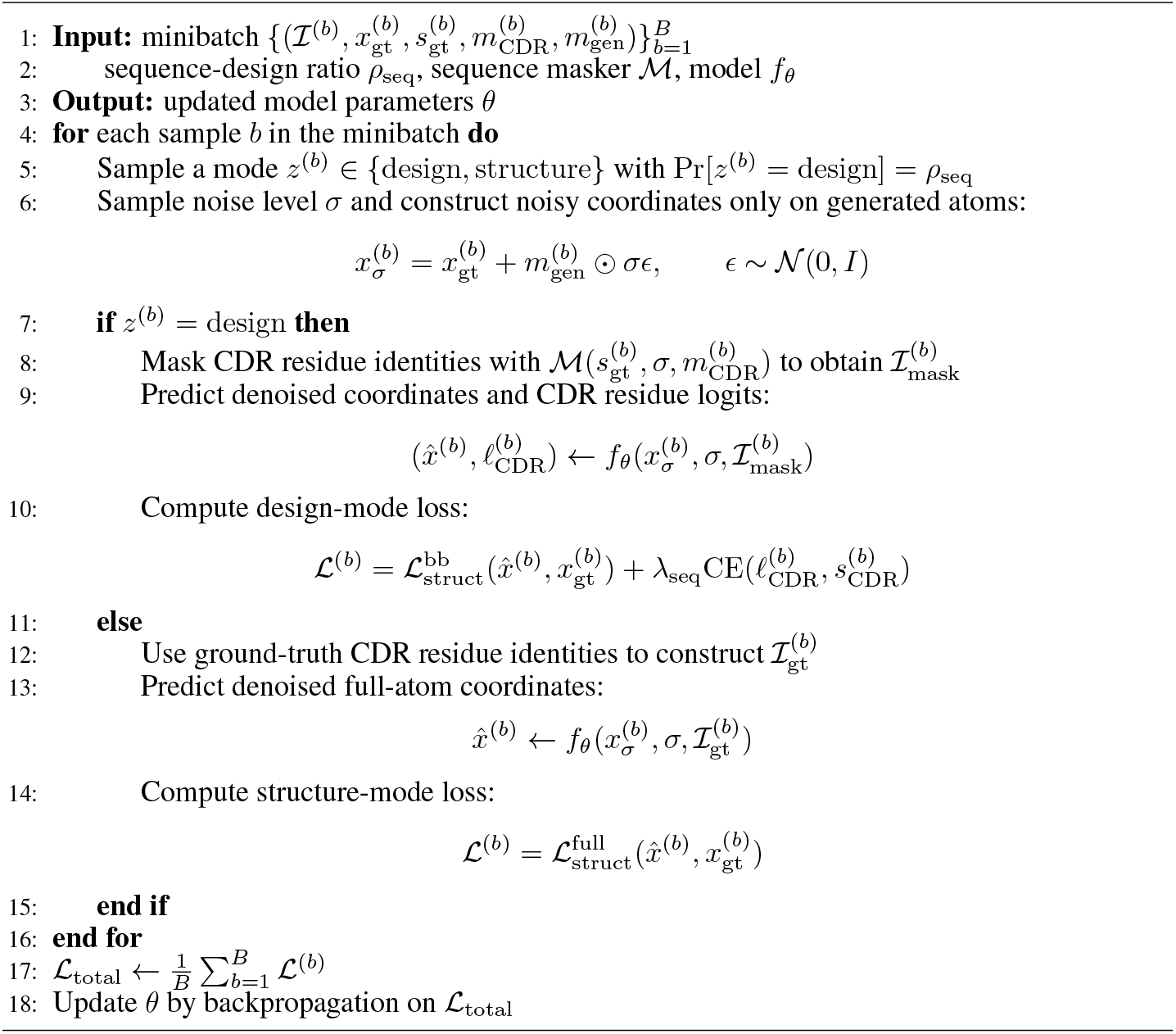

### A Algorithm Details

#### A.1 Mixed Training and Phase-Split Sampling

We provide explicit pseudocode for the mixed training step (Algorithm S1) and the phase-split sampler (Algorithm 1) referenced in Sections 3.3–3.5.

In the implementation, the noise-level threshold σ_switch_ is the only user-facing knob; the equivalent step index T_switch_ used in the main text is computed from σ_switch_ via Algorithm 1, and changes if the number of denoising steps N or the PC strength γ_0_ changes. The training hyperparameter `trunk_cache.sigma_threshold` equals the inference-time σ_switch_, so the phase transition at inference is placed exactly where sequence-design supervision hands over to structure-prediction supervision during training.

#### A.2 Effective SNR Interpretation of the Sequence Mask Schedule

In this appendix, we provide an alternative interpretation of the masking schedule in Eq. (7) through an effective signal-to-noise matching argument.

For coordinate diffusion, the noisy input is

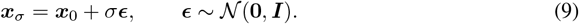

Under the EDM parameterization, the skip coefficient is

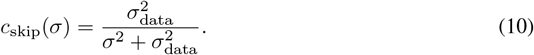

Correspondingly, the coordinate signal-to-noise ratio scales as

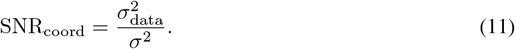

For sequence corruption, let p(σ) be the masking probability. Then the visible fraction is 1 −p(σ) and the masked fraction is p(σ). We define an effective sequence information ratio as

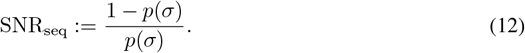

Although this quantity is not a Gaussian SNR in the strict sense, it provides a simple measure of how much visible token information remains relative to the masked portion.

Matching this effective sequence ratio to the coordinate SNR gives

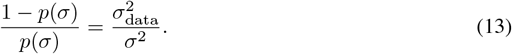

Solving for p(σ) yields

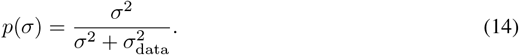

Using the definition of c_skip_(σ), this can be equivalently written as

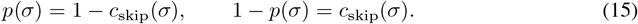

Therefore, the masking schedule adopted in the main text can be understood in two equivalent ways: directly as a visible-fraction alignment rule using the EDM skip coefficient, or as an effective SNR matching rule between sequence and coordinate corruption.

#### A.3 Antibody-Centered Cropping

For antibody/TCR–antigen complexes, the total number of tokens often exceeds the model crop size (e.g., 512 tokens). A naive contiguous or spatial crop, as commonly used in general protein structure prediction, is not ideal for immune complexes: it may discard CDR residues or retain antigen regions that are far from the binding interface. We therefore use an *antibody-centered* cropping strategy tailored to immune complex training (Algorithm S2).

The strategy follows four principles. First, all antibody/TCR tokens are always retained, since they provide the generation target and its structural context. Second, antigen tokens are ranked by distance to the antibody center of mass, so the crop focuses on the interface region. Third, within each antigen chain, selected spans are completed when possible to improve structural continuity. Fourth, during training, the number of antigen tokens is uniformly sampled between a minimum 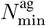 and the maximum allowed by the crop budget, which serves as antigen-context augmentation. At inference time, the crop budget is fixed.

Compared with the mixture of contiguous, spatial, and spatial-interface crops used in general structure prediction, this strategy is deterministic in which chains are preserved and stochastic only in how much antigen context is kept.

#### A.4 Hotspot Conditioning

At training time, epitope residues are computed on the fly from the bound complex and mapped to the model’s is_hotspot_residue input feature. This provides receptor-side interface information and guides generation toward the desired target region.

To avoid over-reliance on fully observed interface annotations, we randomly mask the epitope signal during training. At inference time, the user may provide either the full epitope or a partial subset. The overall procedure is summarized in Algorithm S3.

##### Algorithm S2

Antibody-Centered Cropping

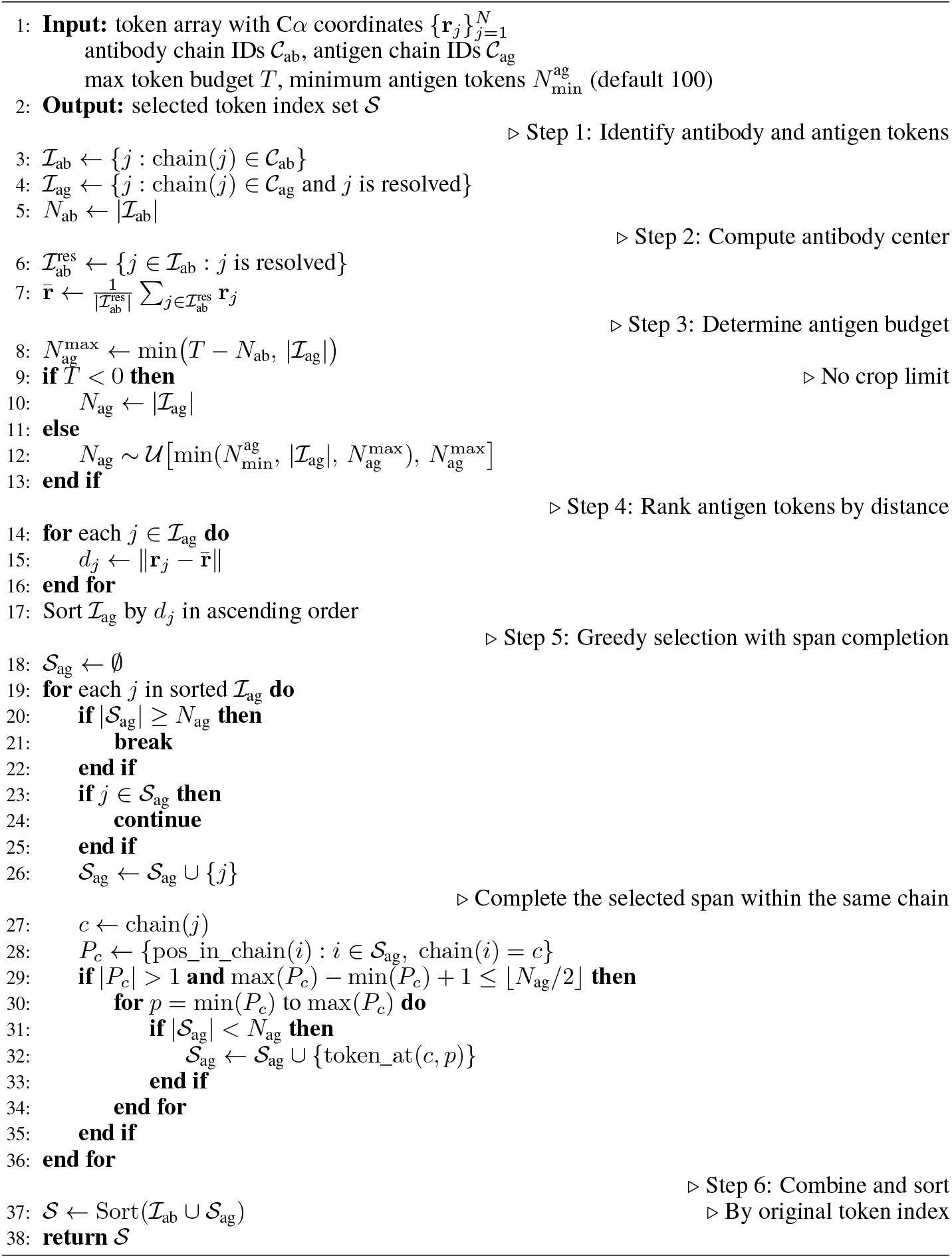

### B Training details and limitations

#### B.1 Training details

All IgGM2 models are trained on 16 A100 GPUs using the atom14 all-atom representation, a max crop size of 512 tokens. We optimize the model with Adam using a learning rate of 1.8 × 10^*−*3^, β_1_ = 0.9, β_2_ = 0.95, and weight decay 1 × 10^*−*8^. The learning rate follows an AF3-style schedule with 3,000 warmup steps, followed by a decay factor of 0.95 every 1,000 steps.

##### Algorithm S3

Epitope Identification and Hotspot Masking

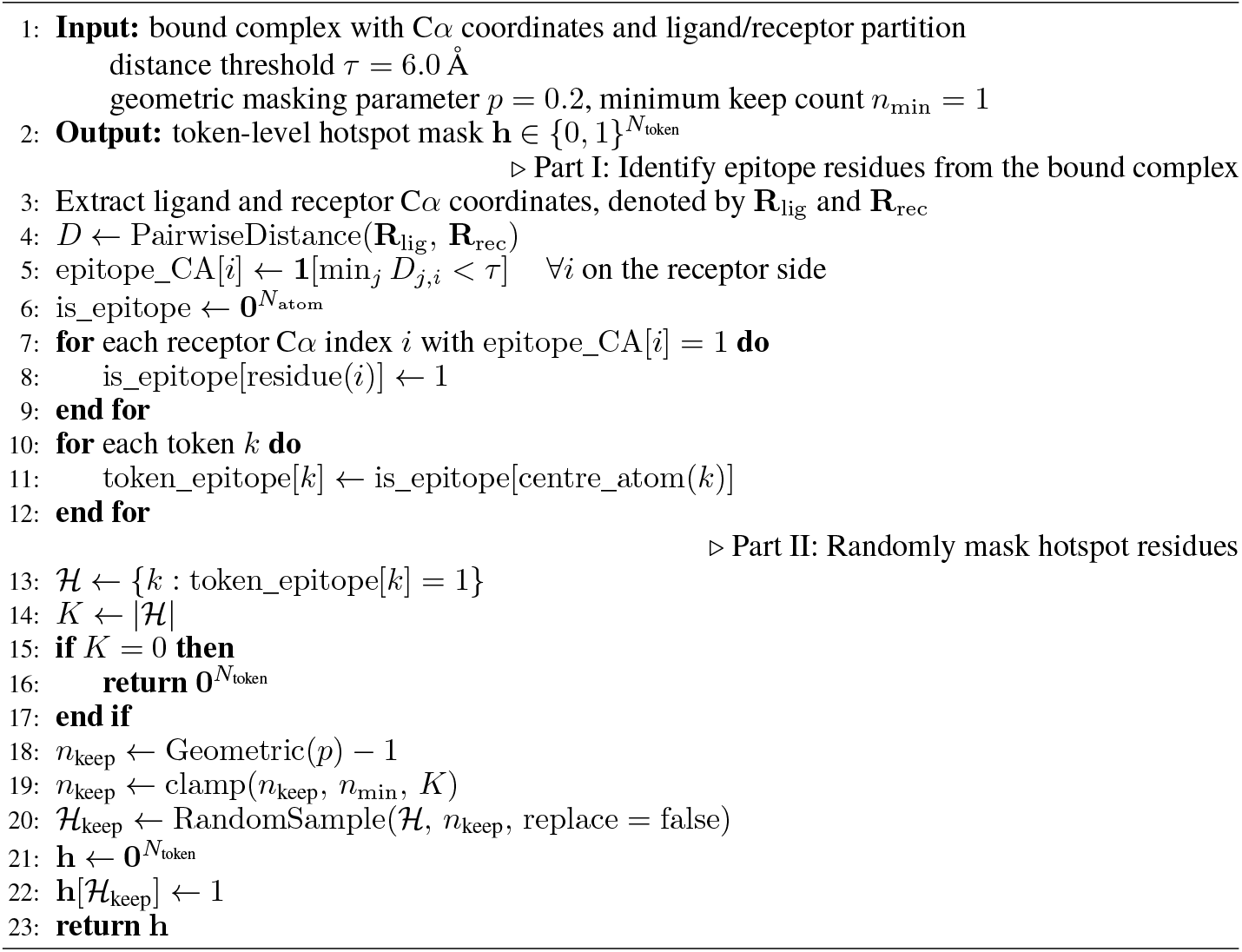

IgGM2-D is initialized from the trained IgGM2-P model and optimized with the same optimizer and learning-rate schedule. We train IgGM2-D with the mixed objective described in Section 3.5, using 30% CDR sequence-design samples and 70% structure-only denoising samples. For sequence-design samples, CDR amino-acid identities are replaced with UNK tokens. We additionally add 0.1Å Gaussian noise to CDR backbone atoms during training.

#### B.2 Limitations

IgGM2 provides a unified full-atom framework for immune receptor structure prediction and CDR sequence–structure co-design, but several limitations remain.

##### Sequence recovery

IgGM2-D still has room to improve amino acid recovery, especially for challenging CDR3 regions and nanobody designs. Although native sequence recovery is not the only goal of de novo design, higher AAR is still useful when preserving native-like binding motifs or functionally important residue patterns. The lower recovery on long HCDR3 loops suggests that stronger sequence priors may help better capture the diversity and constraints of immune receptor repertoires.

##### Staged full-atom refinement

IgGM2-D uses a staged inference procedure, where CDR sequence and backbone geometry are determined before sidechain refinement. This improves sampling stability and avoids separate inverse-folding or external sidechain-packing tools. However, sequence, backbone, and sidechain conformations are tightly coupled, so early errors may affect later sidechain packing. Future work could explore iterative refinement strategies that jointly update sequence, backbone, and sidechains.

##### Dependence on target information

The current design setting assumes that the target structure, and in some cases epitope-level information, is available. This is useful for structure-based design, but performance may depend on the accuracy of the target structure and interface annotations. Predicted antigens, uncertain epitopes, or flexible targets may introduce additional challenges in practical discovery settings.

**Table S1:** IgGM2-P per-subset DockQ / RMSD breakdown on FoldBench under single-seed evaluation.

| Subset | DockQ mean | DockQ median | SR | iRMSD / LRMSD (Å) |
| --- | --- | --- | --- | --- |
| Ab–Ag | 0.611 | 0.751 | 76.23% (93/122) | 3.04 / 8.37 |
| Nano–Ag | 0.574 | 0.762 | 66.67% (32/48) | 4.08 / 9.24 |
| scFv–Ag | 0.843 | 0.843 | 100% (2/2) | 0.85 / 1.66 |
| **Overall** | **0.604** | **0.755** | **73.84% (127/172)** | **3.31 / 8.53** |

| Bin | Range | Count (%) |
| --- | --- | --- |
| High | $\text{DockQ} \geq 0.80$ | 78 (45.3%) |
| Medium | $0.49 \leq \text{DockQ} < 0.80$ | 37 (21.5%) |
| Acceptable | $0.23 \leq \text{DockQ} < 0.49$ | 12 (7.0%) |
| Incorrect | $\text{DockQ} < 0.23$ | 45 (26.2%) |

##### Computational evaluation

Our evaluation mainly relies on computational metrics, including AAR, local Cα RMSD, DockQ, and Rosetta-based interface energy. These metrics capture sequence recovery, structural plausibility, and interface quality, but cannot fully determine binding affinity, specificity, stability, or developability. Experimental validation will be necessary to assess whether generated antibody and TCR designs translate into functional binders.

### C Supplementary Structure Prediction Results

Unless otherwise noted, baseline results in this supplementary structure-prediction section are copied directly from the tFold-TCR paper [Wu et al., 2025b].

#### C.1 FoldBench Per-Subset Breakdown

We report the per-subset breakdown and DockQ-quality distribution behind the main FoldBench result in Table S1. The benchmark contains 172 interfaces from 113 PDB biological assemblies, including 122 Ab–Ag, 48 Nano–Ag, and 2 scFv–Ag interfaces. IgGM2 achieves a mean DockQ of 0.604 on the overall split, with 45.3% of interfaces falling into the high-quality bucket (DockQ ≥ 0.80) and 26.2% into the incorrect bucket (DockQ < 0.23). Unlike the main FoldBench comparison, this analysis provides a more detailed view of performance across receptor types and DockQ-quality levels.

#### C.2 SAbDab-22H2-AbAg Complex Prediction

Table S2 reports the paired antibody–antigen complex prediction results discussed in Section 4.2.

**Table S2:** SAbDab-22H2-AbAg antibody–antigen complex prediction. Unconstrained methods are listed first, followed by methods using contact or epitope restraints.

| Method | Restraint | DockQ ↑ | SR (%) ↑ | TM-Score ↑ |
| --- | --- | --- | --- | --- |
| AlphaFold-3 | – | 0.257 | 32.3 | 0.710 |
| tFold-Ag | – | 0.217 | 28.3 | 0.708 |
| AlphaFold-Multimer | – | 0.158 | 18.2 | 0.665 |
| ColabFold | – | 0.117 | 14.1 | 0.648 |
| tFold-Ag | contact | 0.703 | 97.0 | 0.918 |
| tFold-Ag-ppi | epitope & paratope | 0.416 | 62.6 | 0.814 |
| tFold-Ag-ppi | epitope | 0.303 | 46.5 | 0.761 |
| ColabDock | contact | 0.266 | 41.4 | 0.722 |
| DyMEAN | epitope | 0.160 | 28.3 | 0.700 |
| **IgGM2-P** | epitope | **0.602** | **75.8** | **0.954** |

**Table S3:** SAbDab-22H2-Ab paired-antibody H–L complex prediction. DockQ measures the heavy/light interface quality.

| Method | DockQ ↑ | LRMS ($\text{\AA}$) ↓ | iRMS ($\text{\AA}$) ↓ |
| --- | --- | --- | --- |
| AlphaFold-3 | 0.781 | 1.806 | 1.343 |
| Chai-1 | 0.773 | 1.881 | 1.403 |
| AlphaFold-Multimer | 0.773 | 1.863 | 1.358 |
| tFold-Ab | 0.770 | **1.678** | **1.284** |
| Uni-Fold MuSSE | 0.756 | 1.989 | 1.459 |
| EquiFold | 0.751 | 2.041 | 1.467 |
| ImmuneBuilder | 0.749 | 1.923 | 1.451 |
| DeepAb | 0.722 | 2.079 | 1.564 |
| IgFold | 0.715 | 2.120 | 1.541 |
| **IgGM2-P** | **0.782** | 1.729 | 1.294 |

**Table S4:** SAbDab-22H2-Nano nanobody structure prediction. Per-CDR FR-aligned backbone RMSD (Å, lower is better) under Chothia numbering.

| Method | Fr ↓ | CDR-1 ↓ | CDR-2 ↓ | CDR-3 ↓ |
| --- | --- | --- | --- | --- |
| tFold-Ab | **0.67** | 1.95 | 1.20 | 3.57 |
| AlphaFold-3 | 0.68 | **1.73** | 1.14 | 3.57 |
| Chai-1 | 0.73 | 1.80 | **1.13** | 3.57 |
| AlphaFold | 0.71 | 1.96 | 1.20 | 3.96 |
| ESMFold | 0.70 | 1.87 | 1.34 | 3.80 |
| OmegaFold | 0.73 | 1.77 | 1.34 | 3.63 |
| IgFold | 0.73 | 1.97 | 1.29 | 4.64 |
| ImmuneBuilder | 0.75 | 1.91 | 1.23 | 3.79 |
| HelixFold-Single | 0.76 | 1.92 | 1.27 | 4.16 |
| DeepAb | 0.93 | 2.36 | 1.51 | 9.03 |
| **IgGM2-P** | 0.75 | 2.09 | 1.32 | **3.41** |

**Table S5:** STCRDab-22 unliganded TCR structure prediction (24 samples; 23 αβ-TCR + 1 γδ-TCR). Overall α–β interface docking quality. TCR-specific baselines include tFold-TCR [Wu et al., 2025b] and TCRModel2 [Yin et al., 2023].

| Method | DockQ ↑ | LRMS ($\text{\AA}$) ↓ | iRMS ($\text{\AA}$) ↓ | GDT ↑ |
| --- | --- | --- | --- | --- |
| tFold-TCR | **0.797** | **1.638** | **1.070** | 0.956 |
| TCRModel2 | 0.794 | 1.795 | 1.120 | 0.948 |
| AlphaFold-Multimer | 0.785 | 1.816 | 1.186 | 0.931 |
| AlphaFold-3 | 0.773 | 2.047 | 1.224 | 0.928 |
| **IgGM2-P** | 0.740 | 4.324 | 2.744 | **0.961** |

#### C.3 SAbDab-22H2-Ab

On standalone paired-antibody modeling (Table S3), IgGM2-P achieves the best H–L DockQ among all compared methods, reaching 0.782 and slightly outperforming AlphaFold-3 at 0 781. Across 169 samples, IgGM2-P obtains LRMS of 1.729Å and iRMS of 1.294Å, with a success rate of 100%. It also maintains strong backbone quality, with TM-score 0.951, GDT 0.948, and Cα-RMSD 1.222Å. Its LRMS is second only to the specialized tFold-Ab model, which reaches 1.678Å.

**Table S6:** Co-generated complex quality on the full paired-antibody clean test set. Baselines marked with ^*∗*^ are re-trained on the leak-free training data. HDOCK is used for DiffAb and AbX where external docking is required.

| Method | DockQ ↑ | TM-Score ↑ | CA-RMSD ($\text{\AA}$) ↓ | SR ↑ |
| --- | --- | --- | --- | --- |
| DiffAb* | 0.022 | 0.027 | 3.241 | 0.010 |
| AbX | 0.025 | 0.863 | 1.921 | 0.005 |
| dyMEAN* | 0.087 | 0.864 | 1.425 | 0.082 |
| IgGM | 0.170 | 0.815 | 1.608 | 0.262 |
| RFAntibody | 0.065 | 0.537 | 11.707 | 0.021 |
| **IgGM2-D** | **0.292** | **0.867** | 1.575 | **0.503** |

**Table S7:** Co-generated complex quality on the full nanobody clean test set. Baselines marked with ^*∗*^ are re-trained on the leak-free training data. HDOCK is used for DiffAb where external docking is required.

| Method | DockQ ↑ | TM-Score ↑ | CA-RMSD ($\text{\AA}$) ↓ | SR ↑ |
| --- | --- | --- | --- | --- |
| DiffAb* | 0.023 | 0.006 | 2.146 | 0.000 |
| IgGM | 0.137 | 0.771 | 2.244 | 0.162 |
| RFAntibody | 0.085 | 0.661 | 8.939 | 0.041 |
| **IgGM2-D** | **0.216** | **0.808** | **2.001** | **0.284** |

#### C.4 SAbDab-22H2-Nano

Table S4 reports per-CDR FR-aligned backbone RMSD on the 73-sample nanobody benchmark using Chothia numbering. IgGM2-P achieves the best HCDR3 RMSD, reaching 3.41Å compared with 3.57Å for both tFold-Ab and AlphaFold-3. It also obtains a framework RMSD of 0.75Å and strong overall backbone quality, with TM-score 0.928, GDT 0.919, and Cα-RMSD 1.538Å. Since the input is a single VHH domain without a light chain, interface DockQ is not defined.

#### C.5 STCRDab-22-TCR (unliganded TCR)

Table S5 reports overall α–β interface quality on the 24-sample unliganded TCR benchmark. The benchmark mainly consists of αβ TCRs, with one γδ TCR included following the same chain-mapping convention used by prior baselines. IgGM2-P achieves the best GDT of 0.961, while trailing tFold-TCR [Wu et al., 2025b] and TCRModel2 [Yin et al., 2023] on DockQ and iRMSD. The interface metrics are affected by a small number of outlier cases, but IgGM2-P maintains strong overall backbone quality, with TM-score 0.958, DockQ 0.740, and SR 95.83%.

### D Supplementary Sequence-Design Results

#### D.1 Co-Generated Complex Quality

##### Paired antibody complex quality

Table S6 reports bound-complex quality on the full paired-antibody clean test set by comparing the complexes generated from designed CDR sequences with the ground-truth antibody–antigen complexes. IgGM2-D achieves the best DockQ (0.292), TM-score (0.867), and SR (0.503) among all compared methods. Compared with IgGM, IgGM2-D improves DockQ from 0.170 to 0.292 and nearly doubles SR from 0.262 to 0.503, while maintaining comparable Cα RMSD. These results complement the AAR/RMSD and interface-energy analyses, showing that IgGM2-D better preserves the native antigen-bound geometry after CDR sequence design.

##### Nanobody complex quality

Table S7 reports complex-level quality on the full nanobody clean test set by comparing the co-generated nanobody–antigen complexes with the ground-truth complexes. IgGM2-D achieves the best performance across all reported metrics. Compared with IgGM, it improves DockQ from 0.137 to 0.216 and SR from 0.162 to 0.284, while reducing Cα RMSD to 2.001Å and increasing TM-score to 0.808. These results suggest that the improvements in nanobody AAR and local CDR geometry also translate into better global antigen-bound complex geometry, even in the more challenging long-HCDR3 setting.

**Table S8:**
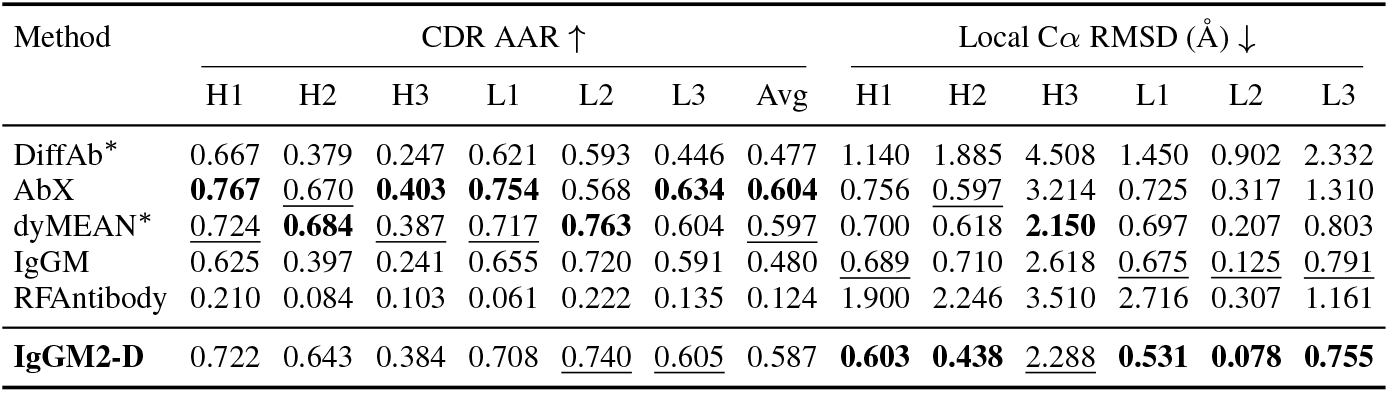
Paired-antibody CDR co-design performance on the full SAbDab design test set before internal test-set deduplication. Higher AAR and lower local Cα RMSD indicate better performance. Best values are bolded and second-best values are underlined. HDOCK is used for DiffAb and AbX when external docking is required.

| Method | CDR AAR ↑ | | | | | | | Local Cα RMSD ($\text{\AA}$) ↓ | | | | | |
| --- | --- | --- | --- | --- | --- | --- | --- | --- | --- | --- | --- | --- | --- |
|  | H1 | H2 | H3 | L1 | L2 | L3 | Avg | H1 | H2 | H3 | L1 | L2 | L3 |
| DiffAb* | 0.667 | 0.379 | 0.247 | 0.621 | 0.593 | 0.446 | 0.477 | 1.140 | 1.885 | 4.508 | 1.450 | 0.902 | 2.332 |
| AbX | **0.767** | <u>0.670</u> | **0.403** | **0.754** | 0.568 | **0.634** | **0.604** | 0.756 | <u>0.597</u> | 3.214 | 0.725 | 0.317 | 1.310 |
| dyMEAN* | <u>0.724</u> | **0.684** | <u>0.387</u> | <u>0.717</u> | **0.763** | 0.604 | <u>0.597</u> | 0.700 | 0.618 | **2.150** | 0.697 | 0.207 | 0.803 |
| IgGM | <u>0.625</u> | 0.397 | <u>0.241</u> | <u>0.655</u> | 0.720 | 0.591 | <u>0.480</u> | <u>0.689</u> | 0.710 | 2.618 | <u>0.675</u> | <u>0.125</u> | <u>0.791</u> |
| RFAntibody | 0.210 | 0.084 | 0.103 | 0.061 | 0.222 | 0.135 | 0.124 | 1.900 | 2.246 | 3.510 | 2.716 | 0.307 | 1.161 |
| **IgGM2-D** | 0.722 | 0.643 | 0.384 | 0.708 | <u>0.740</u> | <u>0.605</u> | 0.587 | **0.603** | **0.438** | <u>2.288</u> | **0.531** | **0.078** | **0.755** |

**Table S9:**
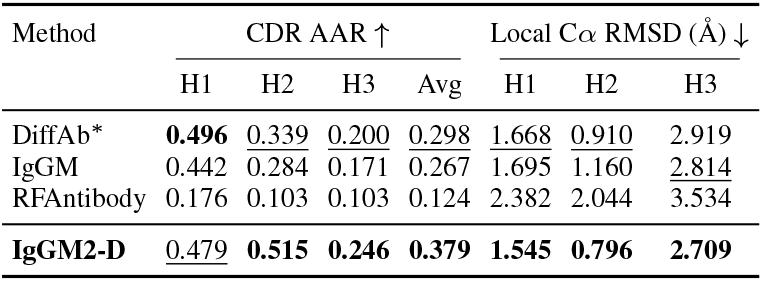
Nanobody CDR co-design performance on the full SAbDab design test set before internal test-set deduplication. AAR is higher better; local Cα RMSD is lower better.

| Method | CDR AAR ↑ | | | | Local Cα RMSD ($\text{\AA}$) ↓ | | |
| --- | --- | --- | --- | --- | --- | --- | --- |
|  | H1 | H2 | H3 | Avg | H1 | H2 | H3 |
| DiffAb* | **0.496** | 0.339 | <u>0.200</u> | 0.298 | 1.668 | 0.910 | 2.919 |
| IgGM | 0.442 | 0.284 | <u>0.171</u> | 0.267 | 1.695 | 1.160 | <u>2.814</u> |
| RFAntibody | 0.176 | 0.103 | 0.103 | 0.124 | 2.382 | 2.044 | 3.534 |
| **IgGM2-D** | <u>0.479</u> | **0.515** | **0.246** | **0.379** | **1.545** | **0.796** | **2.709** |

**Table S10:** Rosetta interface-energy metrics on the full SAbDab design test set before internal test-set deduplication.

| Method | Paired antibodies |  | Nanobodies |  |
| --- | --- | --- | --- | --- |
| | IMP ↑ | Mean $dG$ ↓ | IMP ↑ | Mean $dG$ ↓ |
| DiffAb* | 0.369 | 26.11 | 0.203 | 12.99 |
| AbX | 0.292 | 1.37 | – | – |
| dyMEAN* | 0.314 | 2630.74 | – | – |
| IgGM | <u>0.533</u> | <u>-19.31</u> | <u>0.311</u> | <u>4.86</u> |
| RFAntibody | 0.441 | 94.72 | <u>0.311</u> | 11.74 |
| **IgGM2-D** | **0.595** | **-23.31** | **0.554** | **-24.11** |

**Table S11:** Mean all-six-CDR amino-acid recovery on the TCR interface sequence-design pilot (N=5).

| Method | Designs/target | All CDR | H1 | H2 | H3 | L1 | L2 | L3 |
| --- | --- | --- | --- | --- | --- | --- | --- | --- |
| ProteinMPNN | 10 | 0.433 | 0.518 | 0.350 | 0.446 | 0.390 | 0.400 | 0.450 |
| Rosetta FastDesign | 10 | 0.397 | 0.529 | 0.264 | 0.391 | 0.328 | 0.345 | **0.494** |
| ESM-IF1 | 10 | 0.349 | 0.143 | 0.306 | 0.373 | 0.443 | 0.326 | 0.394 |
| **IgGM2-D** | 1 | **0.626** | **0.840** | **0.833** | **0.466** | **0.633** | **0.807** | 0.455 |

#### D.2 Full-Set Sequence-Design Results

We additionally report CDR co-design results on the full SAbDab design test set before internal test-set deduplication. This full set contains 269 SAbDab design complexes, including 195 paired antibodies and 74 nanobodies. Tables S8, S9, and S10 mirror the main sequence-design tables and report AAR, framework-aligned local Cα RMSD, and Rosetta interface-energy metrics on the full set. The overall trends are consistent with the deduplicated main results, showing that IgGM2-D maintains strong local structural quality and favorable interface-energy metrics even when all test-set sequences are included.

#### D.3 TCR Interface Sequence-Design Pilot

We further evaluate TCR CDR sequence recovery in a small all-six-CDR design pilot with N=5 leak-free TCR-pMHC targets. As shown in Table S11, we compare IgGM2-D with ProteinMPNN, ESM-IF1, and Rosetta FastDesign under the same input complexes and CDR masks. IgGM2-D generates one design per target, while each baseline generates 10 designs per target. We report mean amino-acid recovery over all six TCR CDR loops, as well as region-level AAR for each CDR.

**Table S12:** Full IgGM2-D ablation grid over mixed-training ratio, backbone-noise scale, and phase-transition step. The selected setting is bolded.

| Seq:Struct | Noise | $T_{\text{switch}}$ | All AAR ↑ | Paired AAR ↑ | Nano AAR ↑ | HCDR3 RMSD ↓ | IMP ↑ |
| --- | --- | --- | --- | --- | --- | --- | --- |
| 30/70 | 0.1 | 25 | 0.523 | 0.579 | 0.373 | 3.846 | 0.595 |
| 30/70 | 0.1 | 50 | 0.528 | 0.589 | 0.369 | 3.830 | 0.561 |
| **30/70** | **0.1** | **75** | **0.530** | **0.587** | **0.379** | **3.702** | **0.584** |
| 30/70 | 0.1 | 200 | 0.522 | 0.582 | 0.362 | 3.618 | 0.543 |
| 50/50 | 0.1 | 25 | 0.523 | 0.582 | 0.368 | 3.893 | 0.498 |
| 50/50 | 0.1 | 50 | 0.527 | 0.586 | 0.372 | 3.801 | 0.509 |
| 50/50 | 0.1 | 75 | 0.530 | 0.589 | 0.374 | 3.858 | 0.524 |
| 50/50 | 0.1 | 200 | 0.528 | 0.587 | 0.374 | 3.837 | 0.561 |
| 70/30 | 0.1 | 25 | 0.511 | 0.567 | 0.364 | 3.632 | 0.591 |
| 70/30 | 0.1 | 50 | 0.517 | 0.574 | 0.369 | 3.696 | 0.628 |
| 70/30 | 0.1 | 75 | 0.515 | 0.573 | 0.363 | 3.721 | 0.677 |
| 70/30 | 0.1 | 200 | 0.518 | 0.576 | 0.365 | 3.813 | 0.602 |
| 30/70 | 0.5 | 25 | 0.522 | 0.580 | 0.371 | 3.993 | 0.561 |
| 30/70 | 0.5 | 50 | 0.528 | 0.584 | 0.380 | 4.052 | 0.602 |
| 30/70 | 0.5 | 75 | 0.527 | 0.583 | 0.382 | 4.051 | 0.550 |
| 30/70 | 0.5 | 200 | 0.525 | 0.583 | 0.373 | 3.897 | 0.580 |
| 50/50 | 0.5 | 25 | 0.525 | 0.581 | 0.376 | 3.700 | 0.543 |
| 50/50 | 0.5 | 50 | 0.534 | 0.591 | 0.383 | 3.659 | 0.528 |
| 50/50 | 0.5 | 75 | 0.534 | 0.592 | 0.383 | 3.674 | 0.546 |
| 50/50 | 0.5 | 200 | 0.527 | 0.587 | 0.370 | 3.616 | 0.539 |
| 70/30 | 0.5 | 25 | 0.523 | 0.579 | 0.376 | 3.768 | 0.506 |
| 70/30 | 0.5 | 50 | 0.523 | 0.582 | 0.369 | 3.763 | 0.502 |
| 70/30 | 0.5 | 75 | 0.528 | 0.585 | 0.376 | 3.626 | 0.517 |
| 70/30 | 0.5 | 200 | 0.528 | 0.586 | 0.377 | 3.576 | 0.520 |

### E Supplementary Ablations

#### E.1 Backbone Noise and Full Ablation Grid

The main ablation study identifies two key choices for IgGM2-D: a 30/70 mixed-training ratio and a phase-transition step of T_switch_=75. Table S12 extends this analysis by adding the backbone-noise scale used during sampling, allowing us to examine whether these choices remain robust under different sampling perturbations. We use this full grid to assess the trade-off between sequence recovery and interface quality, rather than selecting the best setting for each individual metric.

Overall, the grid shows that improving AAR alone does not necessarily lead to better designs. For example, increasing the sequence-design fraction or using a larger backbone-noise scale can slightly improve All AAR, with the 50/50 and noise-0.5 setting reaching 0.534. However, these settings reduce IMP compared with our selected configuration. Conversely, the high-IMP 70/30 low-noise setting improves interface preference but lowers All AAR to 0.515. We therefore keep the 30/70 ratio, noise scale 0.1, and T_switch_=75 as a balanced configuration that maintains sequence recovery, nanobody robustness, and interface quality without over-optimizing a single metric.

### F Visualization

#### F.1 TCR–pMHC Complexes

**Figure S1:**
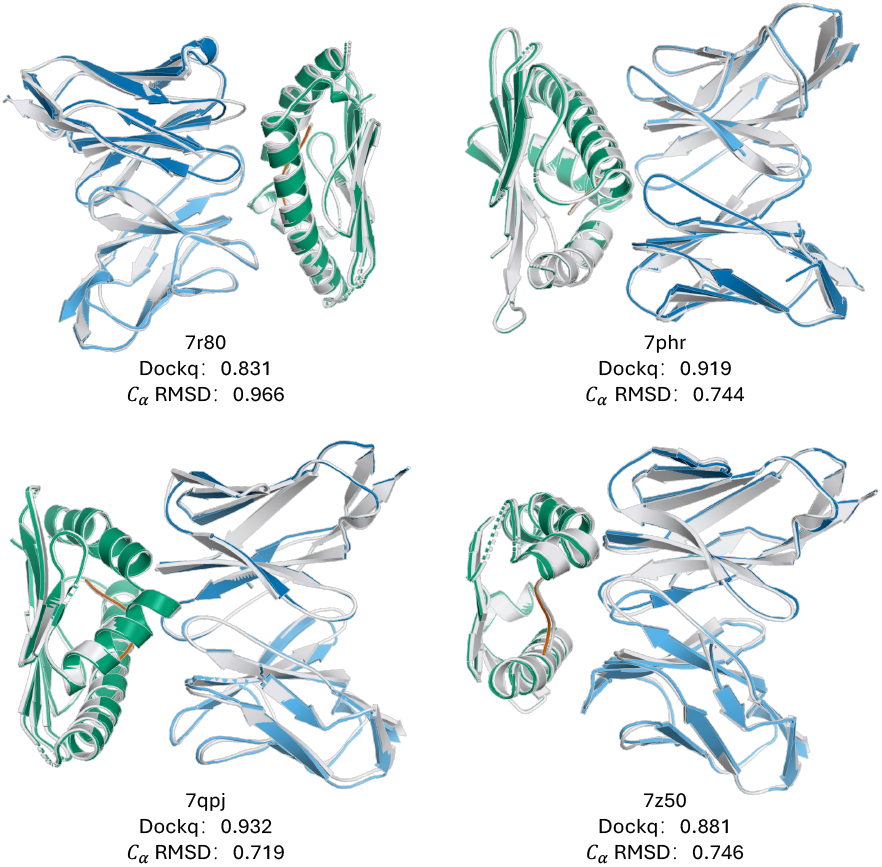
Representative visualization for STCRDab-22-TCR_pMHC complex prediction.

#### F.2 Antibody

**Figure S2:**
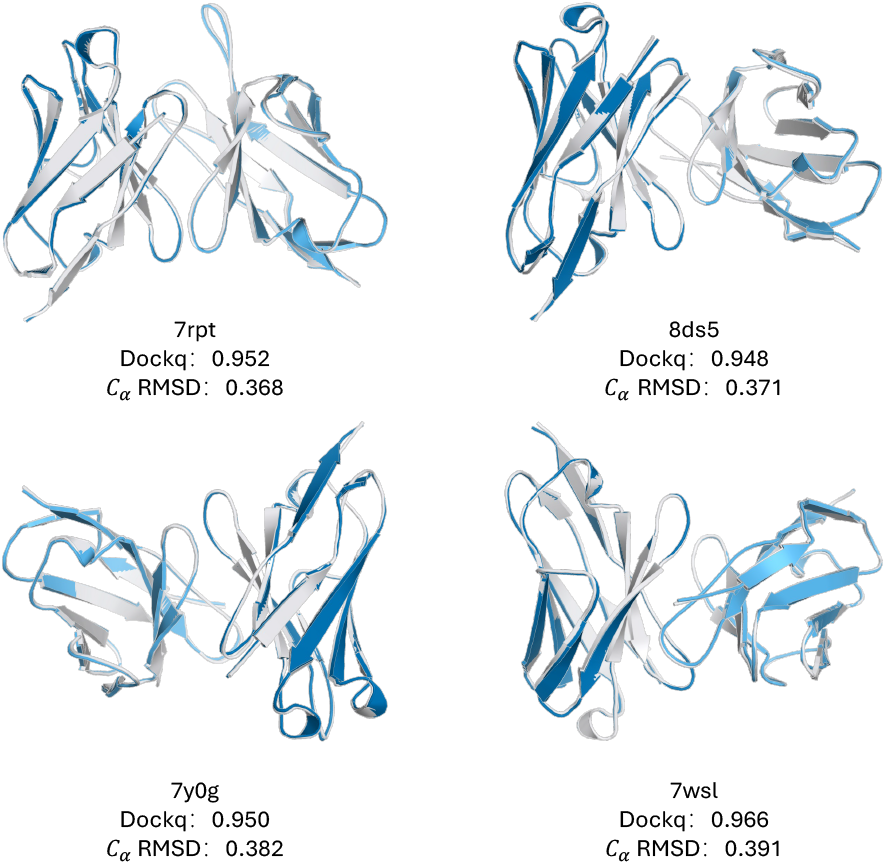
Representative visualization for SAbDab-22H2-Ab antibody prediction.

#### F.3 Nanobody

**Figure S3:**
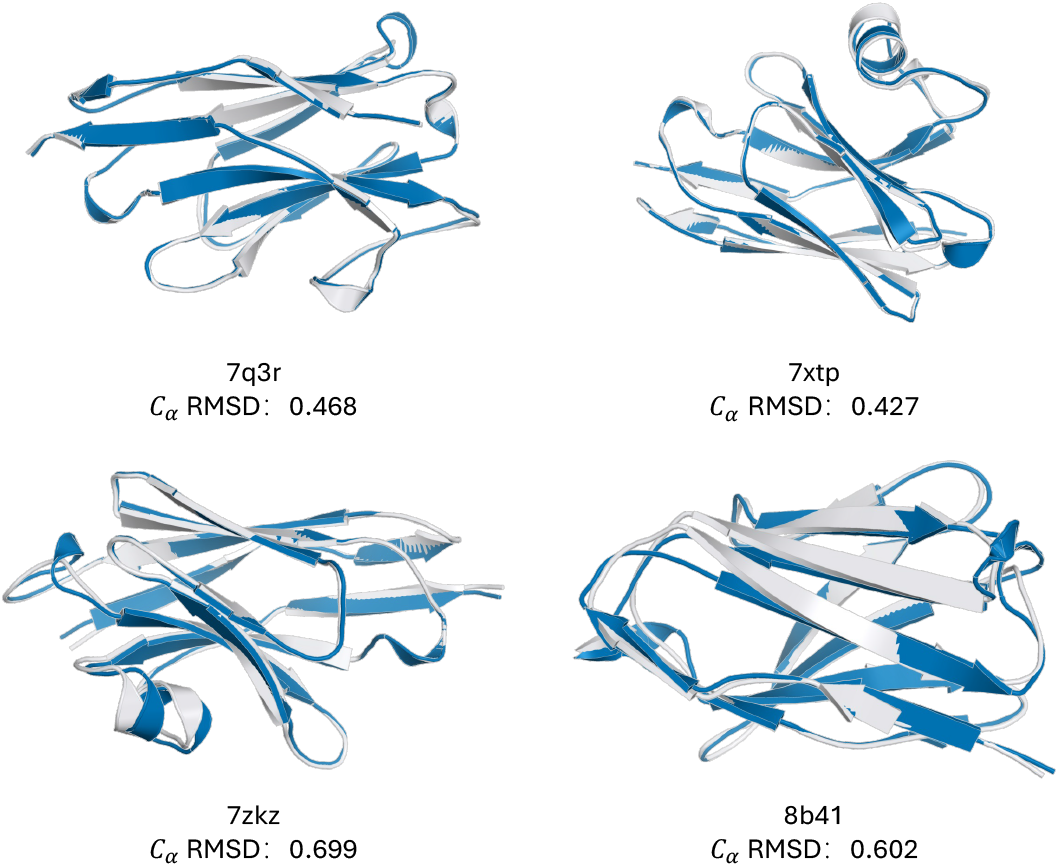
Representative visualization for SAbDab-22H2-Nano nanobody prediction.

#### F.4 Unliganded TCR

**Figure S4:**
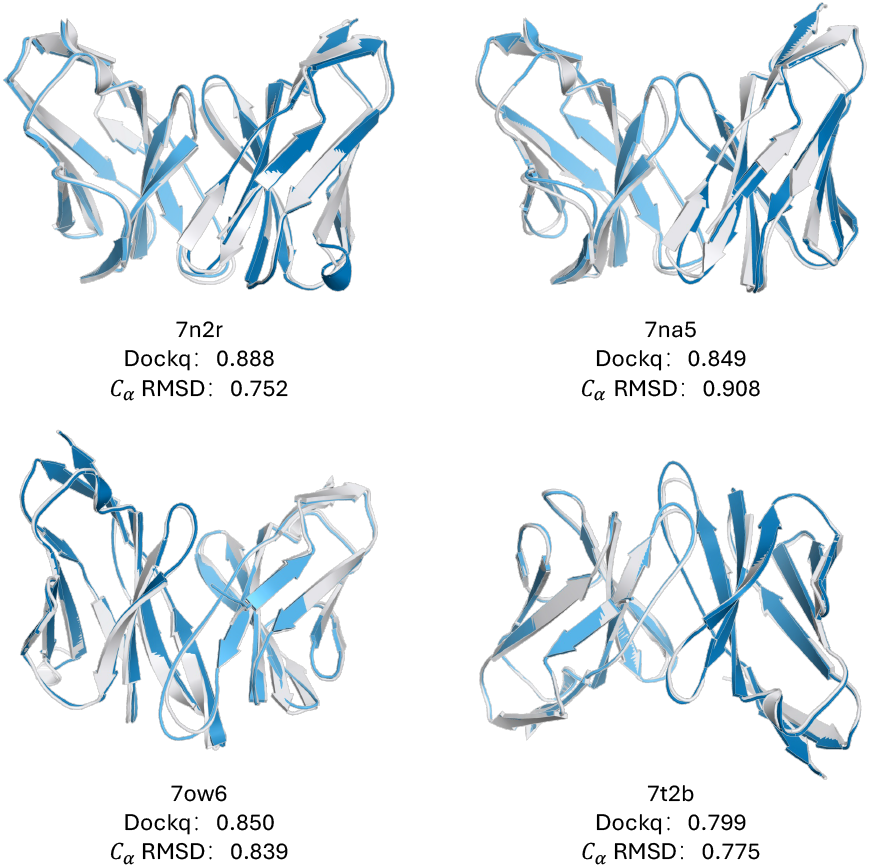
Representative visualization for STCRDab-22 unliganded TCR prediction.

